# Childhood urbanization affects prefrontal cortical responses to trait anxiety and interacts with polygenic risk for depression

**DOI:** 10.1101/246876

**Authors:** Xiao Zhang, Hao Yan, Hao Yu, Xin Zhao, Shefali Shah, Zheng Dong, Guang Yang, Xiaoxi Zhang, Timothy Muse, Jing Li, Sisi Jiang, Jinmin Liao, Yuyanan Zhang, Qiang Chen, Daniel R Weinberger, Weihua Yue, Dai Zhang, Hao Yang Tan

## Abstract

Global increases in urbanization have brought dramatic economic, environmental and social changes. However, less is understood about how these may influence disease-related brain mechanisms underlying epidemiological observations that urban birth and childhoods may increase the risk for neuropsychiatric disorders, including increased social stress and depression. In a genetically homogeneous Han Chinese adult population with divergent urban and rural birth and childhoods, we examined the structural and functional MRI neural correlates of childhood urbanicity, focusing on behavioral traits responding to social status threats, and polygenic risk for depression. Subjects with divergent rural and urban childhoods were similar in adult socioeconomic status and were genetically homogeneous. Urban childhoods, however, were associated with higher trait anxiety-depression. On structural MRI, urban childhoods were associated with relatively reduced medial prefrontal gray matter volumes. Functional medial prefrontal engagement under social status threat during working memory correlated with trait anxiety-depression in subjects with urban childhoods, to a significantly greater extent than in their rural counterparts, implicating an exaggerated physiological response to the threat context. Stress-associated medial prefrontal engagement also interacted with polygenic risk for depression, significantly predicting a differential response in individuals with urban but not rural childhoods. Developmental urbanicity thus differentially influenced medial prefrontal structure and function, at least in part through mechanisms associated with the neural processing of social status threat, trait anxiety, and genetic risk for depression, which may be factors in the association of urbanicity with adult psychopathology.

**Significance Statement:** Urban living has been associated with social inequalities and stress. However, less is understood about the neural underpinnings by which these stressors affect disease risk, and in particular, genetic risk for depression. Leveraging urbanization in China, we studied adults with diverse urban and rural upbringings, who were genetically homogeneous and with similar current socioeconomic status, to isolate the effects of childhood urbanicity. At medial prefrontal cortex, a region critical for processing emotional stressors and social status, genetic risk for depression resulted in more deleterious function under stress in individuals with urban, but not rural childhoods. This implicates medial prefrontal cortex’s critical role in brain development, integrating genetic mechanisms of stress and depression with the childhood environment.

## Introduction

Urban birth and childhoods are associated with risk for schizophrenia(1, 2), autism spectrum disorders(3), substance dependence(4), as well as mood and anxiety disorders(5–7). The world has also been rapidly urbanizing, and in recent years especially so in Asia, bringing dramatic economic, social and environmental changes(8). It is therefore important to better understand how putative environmental and socioeconomic effects in urban versus rural birth and childhoods may influence neural mechanisms of neuropsychiatric risk and resilience(9). Indeed, recent work has implicated an association between urbanicity and social stress in European samples(10–12), but the explicit link to disease susceptibility has been limited.

While the prefrontal cortex and social evaluative stress have been implicated as possible targets of urban exposures(10–12), it is unclear whether and to what extent these targets influence risk-associated enduring trait anxiety-depression, or indeed with genetic risk for disease associated with urban exposures. Studies of medial prefrontal cortex (mPFC) function suggest that this region is critical for incorporating information on social hierarchies in affective regulation(13). Medial PFC updates knowledge about one’s own social position(14), or self-estimates of performance(15), compared to that others, and is critical in navigating complex social relationships and associated stresses. Medial PFC gray matter volume measured on MRI has been associated with strength of interpersonal relationships, social network size and potentially, resilience to stress(16). Reduced mPFC gray matter volumes and altered activation have also been implicated in depression and in associated anxious traits(17–20).

Indeed, if urban birth and childhoods influence mPFC dysfunction through increased sensitivity to social stressors(10, 12), we might expect that these environmental effects would result in structural and functional changes in mPFC associated with states, as well as traits, related to interpersonal stress. Trait anxiety-depression is less studied in this neuroimaging context (but see Lederbogen et al(10)), and is a common association with many psychiatric disorders and their underling genetics(17–21). To test these hypotheses, we examined a genetically relatively homogeneous sample of healthy adult individuals living in Beijing who experienced different levels of rural or urban birth and childhood environments during China’s recent decades of rapid and widespread urbanization, which are amongst history’s largest(22). Based on previous work(23, 24), we designed a novel event-related working memory (WM) task distinguishing between WM maintenance and manipulation under varying social status threat conditions. Because the mPFC is implicated in suppressing self-referential thoughts, especially of social stress in order to execute cognitive tasks such as WM(17, 25–28), we predicted that individuals with urban birth and childhoods, and particularly those with higher trait anxiety-depression, might have more aberrant neural threat responses at mPFC. Critically, we then explore potential gene-environment interactions through which childhood urbanicity may influence stress-related prefrontal responses to polygenic risk for major depressive disorder(29).

## Materials and Methods

### Participants

This study was approved by the Institutional Review Boards of the Peking University Sixth Hospital and the Johns Hopkins University School of Medicine. Following written informed consent, MRI, genetics and questionnaire data were obtained. 522 healthy adults were initially recruited from the local community in Beijing and 490 were included in the current study. (See Supplementary Information for details of inclusion criteria, and clinical and cognitive tests). Of note in this report on social threat and trait anxiety-depression on brain, we used a validated Chinese translation of the Eysenck Personality Questionnaire to study trait anxiety-depression(30, 31), and evaluated cognitive performance using the MATRICS Consensus Cognitive Battery (MCCB)(32–34).

To determine urbanicity, subjects provided residence details from birth to the present. We defined rural areas as agricultural regions with populations typically less than 10,000; urban areas were defined as cities with populations typically more than 100,000 to well over several million. Here, we report results stratifying subjects into an urban group, who had lived in cities since before they were age 12, and a rural group who were born in and lived in rural areas at least until age 12 and beyond. Similar structural and functional MRI results, however, were obtained if we increased the resolution by which differing childhood rural and urban environments are quantified by stratifying subjects into 4 groups, or if we used an urbanicity score based on previous studies(10, 35) (see Supplementary Information).

### Structural MRI data analysis

Subjects were scanned on a 3.0T scanner at the Center for MRI Research, Peking University (see Supplementary Methods for detailed acquisition and analyses procedures). Voxel-based morphometry was performed using SPM8 (http://www.fil.ion.ucl.ac.uk/spm), DPABI(36), and DARTEL(37) to examine urbanicity effects, controlling for age, second polynomial of age, gender, education years, total gray matter volume and MCCB T score(38). Significant effects survived p<0.05 whole-brain family-wise error (FWE) correction.

### Functional MRI task and analysis

We adapted a “number working memory task” based on previous work(23, 24), and included social status threat stressors in half the trials in an event-related block design (Figure 1). Subjects engaged counter-balanced “less stressed” and “stressed” numerical WM maintenance and manipulation trials. Under stress, they performed against a similar age and gender “competitor”, and subsequently received more negative (~70%) than positive feedback about their relative performance. In “less stressed” trials, they engaged without a “competitor” and received neutral feedback. (See Supplementary Information for detailed task parameters.)

**Figure 1.**
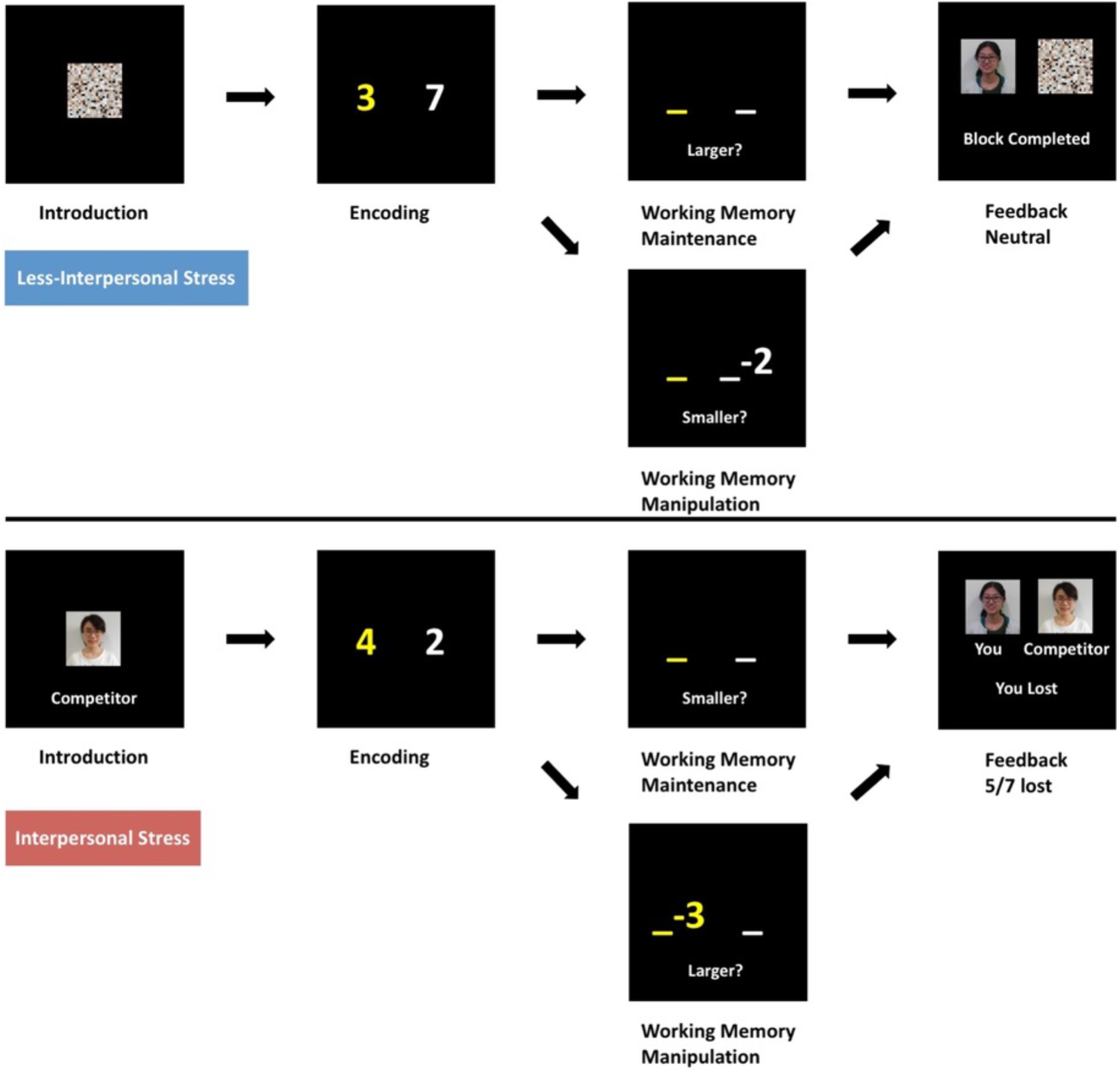
Working memory paradigm incorporating interpersonal threat stress. In the stressed component, subjects were led to believe they were playing against a “competitor” of the similar age and gender, and were judged as winning or losing based on their speed and accuracy, which subsequently resulted in ~70% loss feedback. In the less stressed blocks, there was no competitor and subjects received neutral feedback. In all the working memory manipulation and maintenance tasks, an array of two number digits was encoded and held in working memory over 3-4s. In working memory maintenance, subjects responded to which of the two maintained digits was larger or smaller as indicated. In working memory manipulation, subjects performed subtraction on one of the numbers held in working memory, followed by a response as to which result was larger or smaller as indicated. Subjects performed 2 runs counterbalanced for trial and stimuli presentation order over ~20 mins. All instructions were translated to Chinese.

The functional MRI data were preprocessed and quality-controlled as previously(23, 24), and 394 subjects were included (See Supplementary Information for further details of data preprocessing).

To limit multiple testing and to specifically examine the stress-related function of mPFC, we created 30mm diameter functional ROIs around peaks in the left and right mPFC demonstrably sensitive to stress through the less stress vs stress contrast during WM manipulation, or maintenance, at p<0.05 whole-brain FWE-corrected (See Results, Supplementary Figure S5). From these two ROIs, defined independently of subsequent dependent variables of interest (trait anxiety, urbanicity, or genetic risk for depression), we then examined the hypothesis that trait anxiety-depression is implicated in the expression of states of interpersonal stress at mPFC(20), and how this may be modulated by childhood urbanicity(10). We thus examined the extent to which social threat related engagement at mPFC during WM manipulation might correlate with trait anxiety-depression in each urban and rural sample, and where these effects differed, their potential interaction. We considered significant effects as surviving whole-brain p<0.001 uncorrected and p<0.05 FWE small-volume corrected within the left (542 voxels) or right (722 voxels) mPFC functional ROI. We also performed similar analyses for the WM maintenance task.

As trait anxiety-depression is a risk factor for depressive disorders, also previously associated with urban childhoods(5–7), we then examined the extent to which childhood urbanicity and polygenic risk for depression(29) may influence mPFC function associated with social status threat. Here, we randomly divided our functional MRI dataset into two sub-samples (discovery sample N=200, and replication sample N=194) of approximately equal numbers of individuals with urban or rural childhoods. In the discovery sample, at the same two mPFC ROIs, we examined the extent to which social threat related engagement at mPFC might correlate with MDD polygenic risk score (see below) in each urban and rural group, and where these effects differed, their potential interaction. We considered significant effects as surviving whole-brain p<0.001 uncorrected and p<0.05 FWE small-volume corrected within the mPFC functional ROIs. We then examined similar effects in the replication sample, and in the combined sample.

### Genetic Analyses

DNA collection and genome wide genotyping of the sample is described in Supplementary Methods. Principal Component Analysis (PCA) was performed to determine whether population stratification existed across our urban and rural samples. We evaluated the first 20 PCAs using a two-sample t-test with statistical significance set at p<0.05 corrected for the number of independent components tested.

The polygenic risk score for major depression disorder was based upon a recent genome-wide association study conducted by Psychiatric Genomics Consortium (PGC). The PGC GWAS includes 135,458 cases and 344,901 controls and identify 44 independent loci that were statistically significant(29). Using PLINK v1.07 software(39), we calculated the polygenic risk score based on the 44 lead SNPs of each susceptibility loci. Each SNP was weighted by the effect size from the GWAS, and weighted sums were used to compute the GRS score. Genotype imputation was carried out via the pre-phasing/imputation stepwise approach(40). Genotypes were first phased using SHAPEIT (v2.r727)(41), and imputation was then performed over each 3 Mb interval centered on all index SNPs using IMPUTE (v2.3.0)(40) software. Haplotypes derived from Phase I of the 1000 Genomes Project (release v3) were used as reference data.

## Results

### Demographic and behavioral results

We studied 490 healthy adult subjects with differing birth and childhoods in urban (N=249) and rural (N=241) environments. Subjects with urban birth and childhoods were slightly younger and taller, had higher parental education and parental social-economic status, and older fathers. Rural birth and childhood were associated with a lower total MATRICS Consensus Cognitive Battery (MCCB) cognitive score(32–34) (Table 1). Both groups, however, had similar gender distribution, were all currently living in Beijing, and had achieved similar current educational and occupational levels, controlling for age. Given our hypotheses about social threat and trait anxiety-depression in relation to urbanicity, trait anxiety-depression as measured on the Eysenck Personality Questionnaire Neuroticism sub-score was higher in the urban group (p<0.05, Table 2). Note, however, that none of the subjects were diagnosed with a current or past mood or anxiety disorder.

**Table 1:** Demographic Characteristics of Subjects (N=490)

| | Urban (SD) | Rural (SD) | T/$\chi^2$ | p |
| --- | --- | --- | --- | --- |
| Gender (no. female/male) | 131/118 | 119/122 | 0.512 | 0.474 |
| Age (years) | 23.89 (3.98) | 25.03 (3.45) | 3.389 | 0.001 |
| Education (years) | 16.63 (2.32) | 16.98 (2.62) | 1.579 | 0.115 |
| Height (cm) | 169.27 (8.13) | 167.65 (7.61) | 2.265 | 0.024 |
| Weight (kg) | 62.42 (13.04) | 61.35 (11.49) | 0.961 | 0.337 |
| Father's Childbearing Age (years) | 28.45 (4.74) | 27.02 (6.00) | 2.909 | 0.004 |
| Father's Education (years) | 12.94 (3.57) | 9.59 (3.29) | 10.805 | <0.001 |
| Mother's Childbearing Age (years) | 26.63 (3.83) | 25.92 (5.08) | 1.746 | 0.081 |
| Mother's Education (years) | 12.42 (3.55) | 7.88 (3.88) | 13.515 | <0.001 |
| Current Occupation (no.) |  |  | 3.724 | 0.590 |
| Managers in public or private companies: Urban (5), Rural (5); |  |  |  |  |
| Professionals and Technicians: Urban (46), Rural (58); |  |  |  |  |
| Clerks: Urban (12), Rural (10); |  |  |  |  |
| Service Personnel: Urban (13), Rural (10); |  |  |  |  |
| Students: Urban (164), Rural (145); |  |  |  |  |
| Others: Urban (9), Rural (13); |  |  |  |  |
| MCCB | 54.14 (4.43) | 51.03 (4.89) | 7.380 | <0.001 |
| Total GMV (in cm<sup>3</sup>) | 752.13 (61.89) | 739.12 (61.58) | 2.334 | 0.020 |

GMV, Grey Matter Volume. MCCB: MATRICS Consensus Cognitive Battery

**Table 2:** Demographic Characteristics of Subjects in the Functional MRI Study (N=394)

| | Urban (SD) | Rural (SD) | T/$\chi^2$ | p |
| --- | --- | --- | --- | --- |
| Gender (female/male no.) | 109/90 | 99/96 | 0.634 | 0.426 |
| Age (years) | 23.69 (3.65) | 24.96 (3.13) | 3.707 | <0.001 |
| Education (years) | 16.63 (2.32) | 17.20 (2.53) | 2.337 | 0.020 |
| EPQ Neuroticism | 7.60 (4.88) | 6.65 (4.66) | 1.968 | 0.0498 |
| **Stress** |  |  |  |  |
| Accuracy in WM manipulation | 0.86 (0.10) | 0.86 (0.10) | 0.471 | 0.638 |
| RT in WM manipulation (s) | 1.43 (0.34) | 1.54 (0.36) | 2.901 | 0.004 |
| Accuracy in WM maintenance | 0.92 (0.08) | 0.92 (0.10) | 0.903 | 0.367 |
| RT in WM maintenance (s) | 1.04 (0.27) | 1.11 (0.27) | 2.660 | 0.008 |
| **Less Stress** |  |  |  |  |
| Accuracy in WM manipulation | 0.85 (0.10) | 0.86 (0.10) | 0.650 | 0.516 |
| RT in WM manipulation (s) | 1.48 (0.34) | 1.58 (0.36) | 2.789 | 0.006 |
| Accuracy in WM maintenance | 0.86 (0.08) | 0.86 (0.09) | 0.442 | 0.659 |
| RT in WM maintenance (s) | 1.10 (0.31) | 1.19 (0.31) | 3.010 | 0.003 |

Abbreviations: EPQ, Eysenck Personality Questionnaire; RT, Reaction Time; WM, Working Memory.

The Han Chinese subjects were genetically homogeneous with no significant differences across the first 20 principal components from genome-wide genotyping (Supplementary Figure S1). Increasing the resolution by which childhood urban-rural exposures are defined, using four groups or calculating an urbanicity score(10, 35), yielded similar demographic, behavioral, genetic and MRI structural and functional findings described below (see Supplementary Methods, Supplementary Figure S3, S4, S8).

During WM maintenance, trials with status threat were associated with relatively increased accuracy compared with less stress (p<0.001, Table 2). There was, however, no stress-related increased accuracy in WM manipulation, resulting in a significant task-by-stress interaction (F=33.1, p<0.001), consistent with a well-established bias for perseverative as opposed to flexible WM operations under stress(42) (Supplementary Figure S2, Supplementary Table S1). During WM maintenance and manipulation trials, the evoked stress was associated with faster reaction times, though without a task-by-stress interaction (Supplementary Table S1).

Across urban vs rural childhoods (Table 2), there were no differences in accuracy, but relatively faster reaction time in the urban vs rural groups in WM manipulation, as well as in WM maintenance, whether during the stressed, or less-stressed conditions (p<0.01). There was, however, no urbanicity-by-stress interaction on accuracy, or on reaction time during WM manipulation, or WM maintenance. Thus, while there were no behavioral confounders for several of the key urbanicity-by-stress related neuroimaging contrasts (see below), possible occult demographic and behavioral effects were nevertheless controlled for in functional MRI analyses through the event-related design matrix focusing on correct trials, as well as the inclusion of reaction time and age as additional covariates.

### Effects of urban vs rural childhoods on gray matter volume

Total gray matter volume was greater in individuals in the urban compared with rural childhoods (p=0.02). Despite this global effect, when controlled for age, the second-degree polynomial expansion of age, gender, education, total gray matter volume and MCCB T score, individuals with rural childhoods had significantly increased regional gray matter volume in the mPFC (p<0.05, whole-brain family-wise-error, FWE, corrected, Figure 2) in Brodmann Area (BA) 11 (x=−6, y=59, z=−20; T=4.81) and BA8 (x=8, y=33, z=40; T=4.73; p<0.05 whole-brain FWE corrected). No other brain regions differed between childhoods in urban and rural environments at these thresholds. As mPFC is critically engaged in processing hierarchical social threat stress(13, 14, 43) potentially mediating urban-rural effects on brain, we then focused on this region-of-interest (ROI) in subsequent functional MRI analyses below.

**Figure 2.**
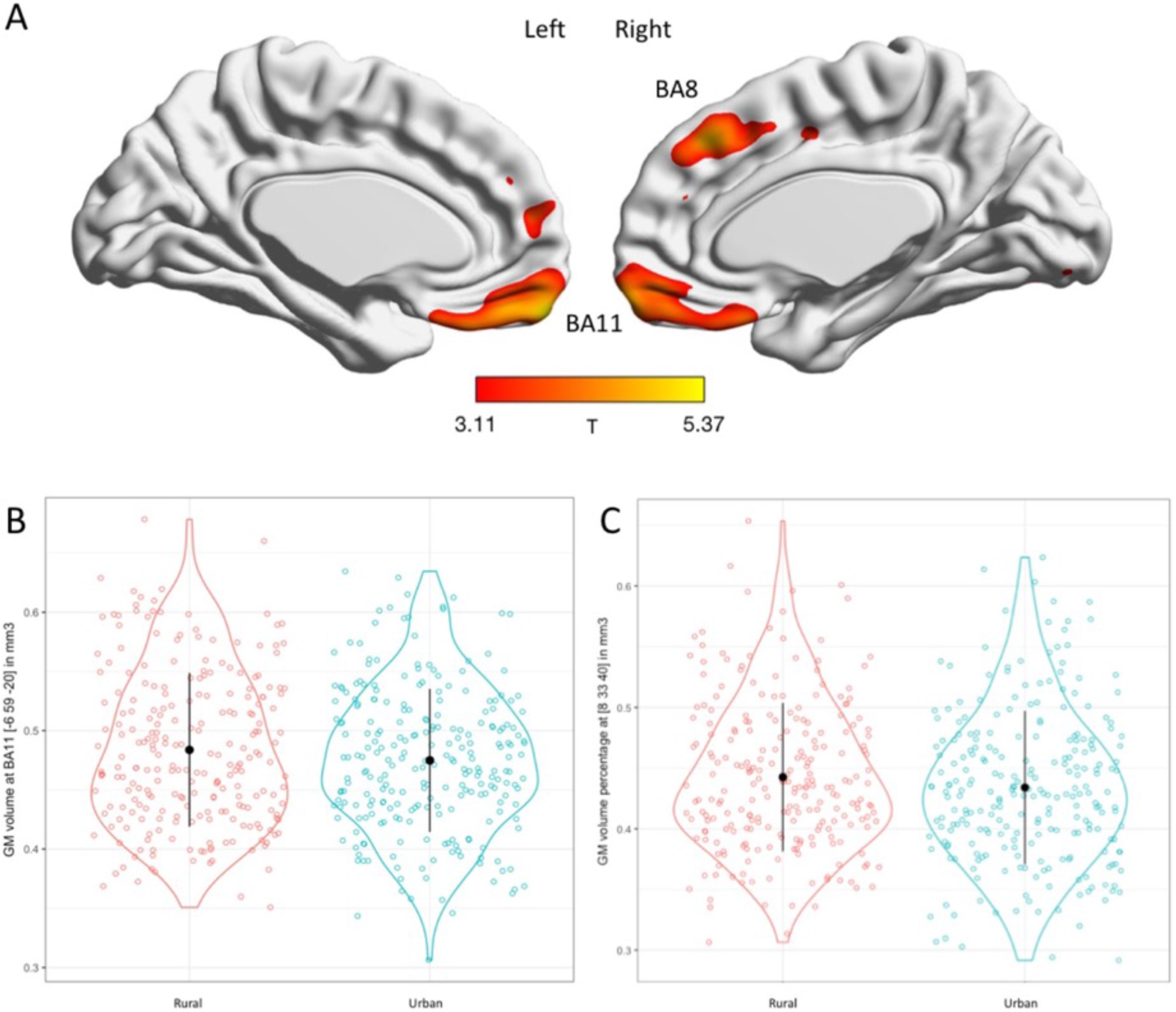
Childhood urbanicity effect on grey matter volume (N=490) (A) Brain map of the rural vs urban gray matter volume effects controlled for age, second polynomial of age, gender, education, total grey matter volume, MCCB T score (shown at p<0.001 uncorrected but peaks survived p<0.05 whole-brain FWE corrected). (B) Plot of rural vs urban effects at Brodmann area 11 in medial prefrontal cortex (peak x=−6 y=59 z=−20, t = 4.81, p<0.05, whole brain FWE corrected). (C) Plot of rural vs urban effects at Brodmann area 8 in mPFC (peak x=8 y=33 z=40, t = 4.73, p<0.05, whole-brain FWE corrected).

### Effects of social status threat, trait anxiety and urbanicity on medial PFC function

During each of the WM maintenance and manipulation tasks, regions in the prefrontal, parietal and temporal cortices, and striatum, were robustly engaged, along with well-established decreased engagement of the mPFC, putatively associated with the degree to which self-representations needed to be suppressed to perform active cognitive tasks(17, 25–28) (p<0.05 whole-brain FWE-corrected, Supplementary Figure S5, Supplementary Table S2). Relative to maintenance, WM manipulation engaged greater activity at the prefrontal cortex, parietal lobe, temporal lobe, and striatum, as well as more suppression of the mPFC (Supplementary Table S3). Under putatively downgraded social status in the stressed vs the less stressed condition, there was greater suppression of mPFC engagement during WM manipulation, as well as in WM maintenance (Supplementary Figure S5, Supplementary Table S4). Medial PFC suppression was correlated with relatively poorer indices of performance in reaction time (p<0.05 whole brain FWE corrected, Supplementary Figure S6) and lower overall cognitive MCCB score (Supplementary Figure S7). Accuracy was not significantly correlated with brain activation, consistent with the event-related design considering only correctly performed trials. We then focus specifically on the stress-related function of mPFC, where we created 30mm diameter functional ROIs around peaks in the left and right mPFC demonstrably sensitive to stress through the less stress vs stress contrast during WM manipulation, or maintenance, at p<0.05 whole-brain FWE-corrected (Supplementary Figure S5).

As trait anxiety-depression, which was higher in those with urban childhoods (Results above), could potentiate the effects of social status threat(44, 45), we then examined how this trait modulated stress-related mPFC function in individuals with urban childhoods, and how this compared with rural childhoods in the two mPFC ROIs. In individuals with urban childhoods, higher trait anxiety-depression was associated with greater suppression of mPFC engagement during WM manipulation under social threat stress (Figure 3A, x=18, y=60, z=6, T=3.26, p<0.001 uncorrected, p<0.05 FWE small-volume corrected within the mPFC ROI). On the other hand, in those with rural childhoods, these effects were not significant at these thresholds. This resulted in a significant interaction at the mPFC in which subjects with urban childhoods and higher trait-anxiety-depression required apparently greater suppression of self-related mPFC function(17, 25–28) to perform WM manipulation under social threat stress (Figure 3B, x=8, y=54, z=−4, T=3.37, p<0.001 uncorrected, p<0.05 FWE small-volume corrected within the mPFC ROI). Effects during WM maintenance were not significant at these thresholds (Supplementary Figure S9). The results are unlikely to be confounded by task performance relationships with trait anxiety-depression as there were no significant correlations between trait anxiety-depression and accuracy or reaction time during WM maintenance or manipulation; there was also no urbanicity-by-accuracy or urbanicity-by-reaction time interaction for WM maintenance, or for WM manipulation.

**Figure 3.**
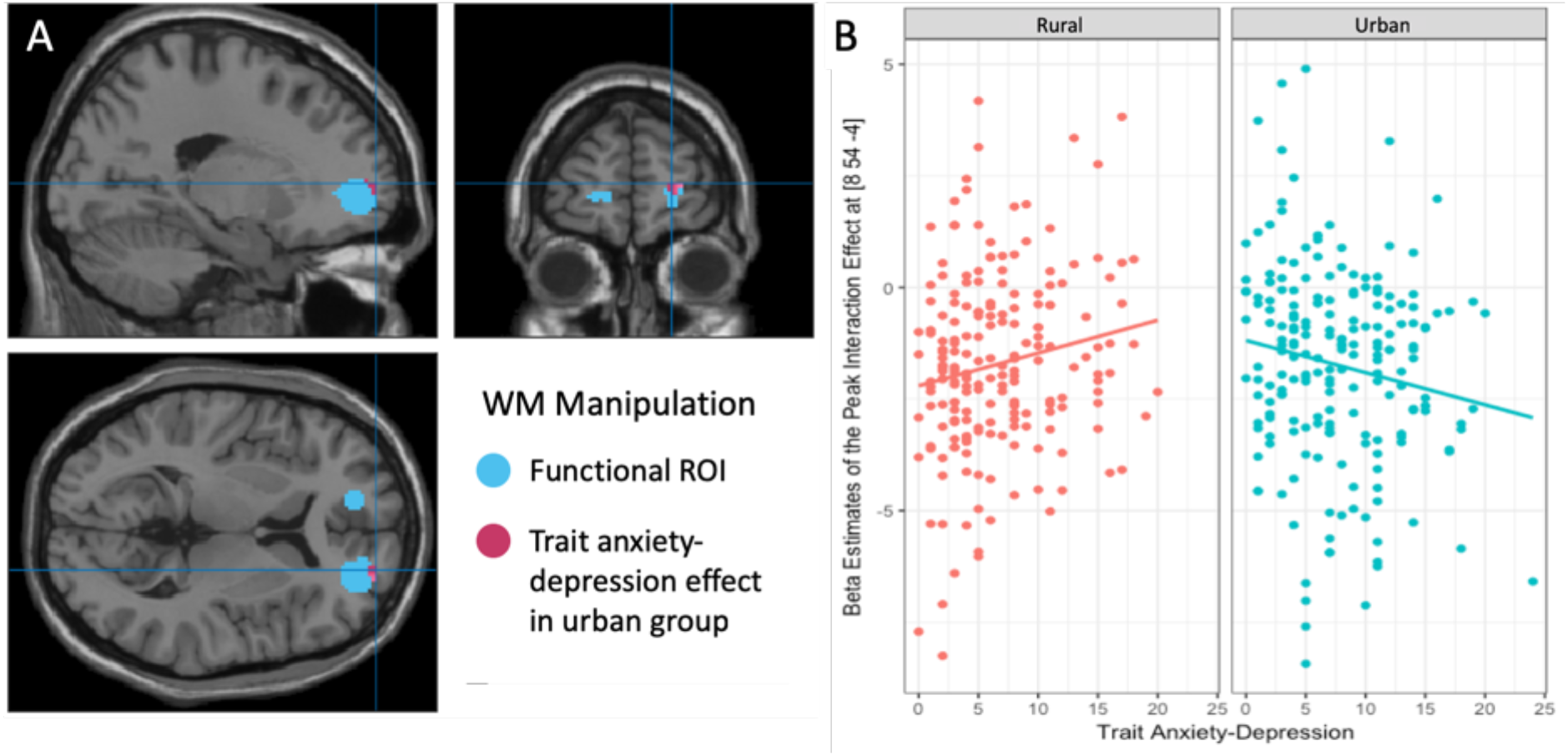
Effects of childhood urbanicity on medial PFC function during working memory manipulation under social threat stress. (A) Brain map showing the mPFC functional ROIs (blue), and peak correlation of mPFC suppression with increasing trait anxiety-depression during the WM manipulation task under social threat in individuals with urban childhoods (N=199, x=18, y=60, z=6, T=3.26; p<0.001, p<0.05 FWE corrected within the right functional ROI). (B) Plot of the interaction between rural and urban childhoods, and trait anxiety-depression on mPFC engagement (N=394, x=8, y=54, z=−4, T=3.37; p<0.001 uncorrected, p<0.05 FWE corrected within the functional ROI). In the group with more urban childhoods, increased trait anxiety-depression was associated with in a larger reduction in mPFC engagement during stress. In subjects with rural childhoods, however, this effect was less apparent.

### Interaction of urbanicity and polygenic risk of major depressive disorder on social threat-related medial PFC function

In the discovery (N=200 with 104 urban and 96 rural childhoods), replication (N=194 with 99 urban and 95 rural childhoods) and combined samples, polygenic risk score for depression did not differ across urban and rural childhoods, and was not associated with demographic variables, or with WM task accuracy or reaction time under stress, or less stress. In the each sample, at the same two mPFC ROIs, we then examined the extent to which social threat related engagement at mPFC might correlate with MDD polygenic risk score in each urban and rural group, and where these effects differed, their potential interaction. In the discovery sample, subjects with urban childhoods and higher polygenic risk for depression had greater suppression of mPFC engagement under social threat stress relative to less stress during WM manipulation (Figure 4A; x=−18 y=54 z=02, T=4.47, p<0.001 uncorrected, p<0.05 FWE small-volume corrected within the mPFC ROI). These effects were not significant in subjects with rural childhoods, resulting in a gene-environment interaction at the mPFC (Figure 4B; x=−18 y=54 z=−6, T=3.58, p<0.001 uncorrected, p<0.05 FWE small-volume corrected within the mPFC ROI).

**Figure 4.**
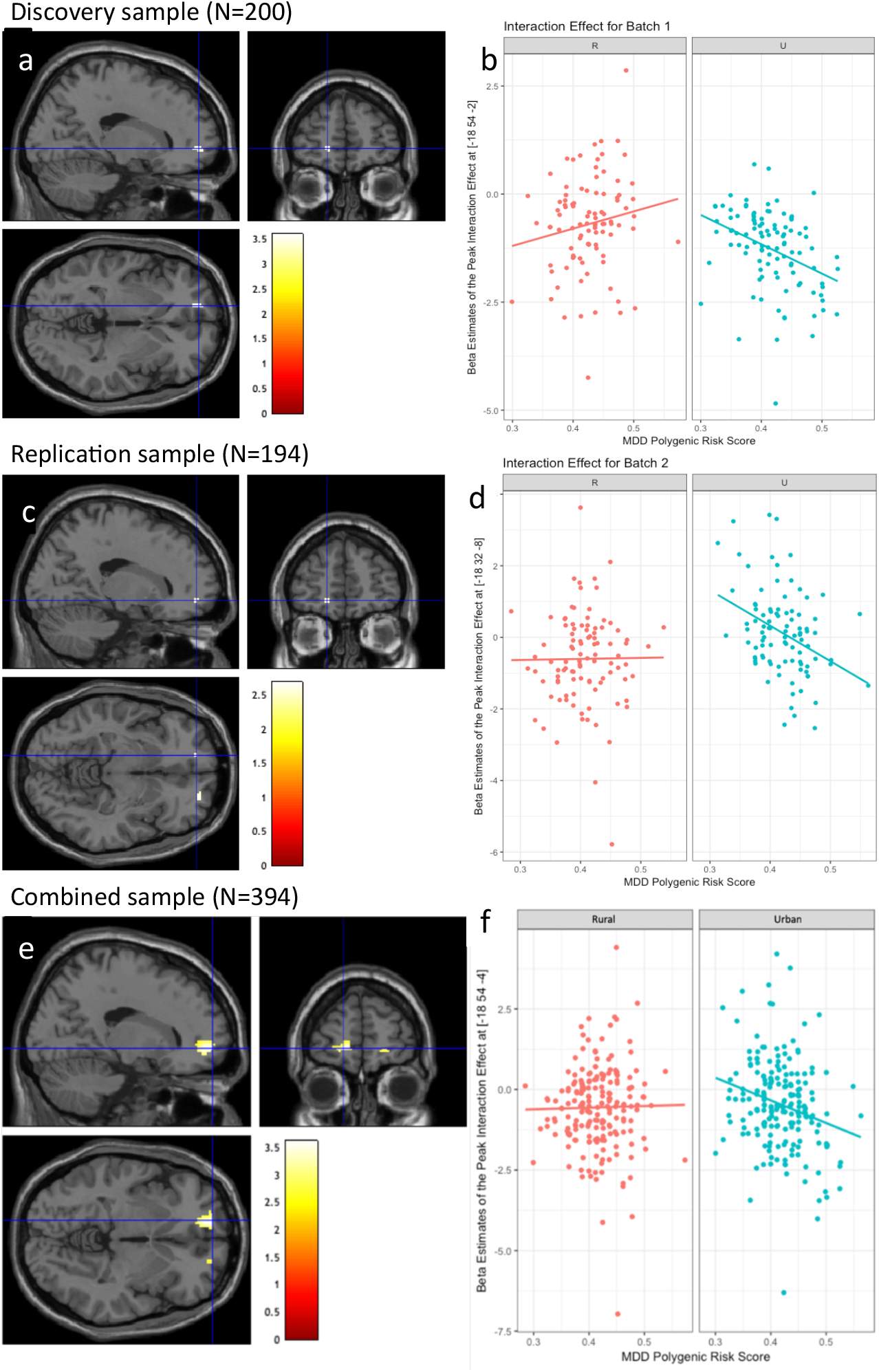
Effects of childhood urbanicity and polygenic risk for depression on stress-associated mPFC function during working memory manipulation. (A) In the discovery sample, peak effects in subjects with urban childhoods where polygenic risk for depression correlated with relatively deleterious reduced engagement of mPFC under stress vs less stress (p<0.05 FWE small volume corrected within the functional ROI). (B) Plot of the gene-environment interaction between rural or urban childhoods and polygenic risk for depression on stress-related mPFC engagement, p<0.05 FWE small volume corrected within the functional ROI). In the group with urban childhoods, increased polygenic risk for depression resulted in reduced medial prefrontal cortex engagement during stress. This effect was less apparent in those with rural childhoods. (C) Similar effects were observed in the replication sample, where those with urban childhoods had more deleterious engagement of stress-related mPFC in relation to polygenic risk for depression. (D) These effects were absent in those with rural childhoods, resulting in a gene-environment interaction (p<0.005 uncorrected). (E) In the combined sample, there was a correlation of polygenic risk for depression with deleterious mPFC engagement (p<0.05 FWE small-volume corrected) in those with urban childhoods. (F) The absence of these effects in those with rural childhoods resulted in a significant gene-environment interaction (p<0.05 FWE small-volume corrected) in the combined sample. All brain maps shown with thresholding at p<0.001 uncorrected.

In the replication sample, subjects with urban childhoods also had more suppressed mPFC engagement during WM manipulation under social stress relative to less stress that correlated with polygenic risk for depression (Figure 4C; x=−14 y=56 z=−6, T=4.46, p<0.001 uncorrected, p<0.05 FWE small-volume corrected). These effects were also not significant in subjects with rural childhoods, resulting in a gene-environment interaction at the mPFC (Figure 4D; x=−18 y=62 z=−4, T=2.54, p<0.005 uncorrected). In the combined sample, the effect of stress-related mPFC engagement on polygenic risk for depression in the subjects with urban childhoods (Figure 4E; x=−18 y=58 z=−4, T=3.61, p<0.001 uncorrected, p<0.05 FWE small-volume corrected within the mPFC ROI), but not those with rural childhoods, resulted in a gene-environment interaction at the mPFC (Figure 4F; x=−28 y=58 z=−2, T=3.23, p<0.001 uncorrected, p<0.05 FWE small-volume corrected within the mPFC ROI). These effects were not present at the mPFC ROIs during stress-related WM maintenance.

## Discussion

We examined the effects of urban and rural birth and childhoods on brain structure and social status threat related brain function in a large and genetically homogeneous sample of healthy adult Han Chinese individuals. The sample is unique in terms of having had similar current educational and occupational status, and genetic ancestry, but different childhoods during China’s recent rapid urbanization. This has allowed us to isolate the role of urban vs rural childhoods on brain development and function in the context of genetic risk for depression that have previously not been attainable. Prior work has highlighted the role of mPFC in mediating increased social stress sensitivity and urbanicity in European populations(10), but these samples have not been genetically controlled and the association with risk-related psychological traits and illness associated genetic risk has been unclear. We find that urban birth and childhoods affect the MRI structure and function of mPFC, the latter more specifically associated with processing active WM tasks under social threat stress, in which individuals with urban childhoods and higher trait anxiety-depression had apparently more “deleterious” function. Furthermore, the stress-related effects at mPFC were correlated with polygenic risk score for major depressive disorder in association with urban but not rural childhoods, in a significant gene-environment interaction. These data implicate childhood urbanicity in mediating the genetic mechanisms of depressive illness on mPFC function.

Our findings of reduced mPFC gray matter volumes in relation to urban birth and childhoods may indirectly implicate increased adverse stress exposure. Medial PFC gray matter volume reductions have been observed in childhood adversity(46, 47), depression(19), post-traumatic stress disorder(48), and in relation to increased trait anxiety(20). We note, however, that none of our subjects suffered from a psychiatric illness or had evidence of childhood maltreatment, and gray matter volume reductions in our neurotypical sample are relative in association with the early childhood environment. Our mPFC findings are consistent with that in a study from Germany, where urban childhoods were associated with significantly reduced lateral and medial prefrontal gray matter volumes, though their results were more robust in males(11). The lateral and medial prefrontal changes are part of a similar network processing adverse experiences(49), particularly given the cognitive appraisal of these emotional experiences(50). While we did not detect gender dimorphic results, these could reflect cultural dissimilarities, or our larger sample size. Ours is also the first study of childhood urbanicity with genomic control.

The underlying mechanisms by which urban birth and childhood is associated with prefrontal cortical variation is unknown. There is prior evidence that urban childhoods may be characterized by more stressful social environments, greater socioeconomic disparities and associated threats to social status(51–53), perhaps more so in China(54). Parental stress may also affect pre and postnatal child development(55), and have been associated with increased trait anxiety-depression(56–58). These enduring behavioral traits may then contribute to states of stress through tendencies to mis-appraise the social environment as threatening and coping resources as low(59–62). Cognitive appraisal engages executive function and lateral and medial prefrontal cortex(49), also known to be sensitive to stress, at least in part through dopaminergic mechanisms(63). Indeed, maladaptive self-referential ruminations characteristic of trait anxiety-depression(64) relate to mPFC function, which under active cognitive task demands is suppressed(17, 25–28). These prior studies suggest that if indeed urban birth and childhoods affect cortical function under stress, they would impact enduring traits related to the processing of stress, and executive function under stress. These assumptions are consistent with our findings, where urban childhoods potentiated the effect of even normal variation in trait anxiety-depression to greater extent in mPFC function under social and cognitive stress during WM manipulation.

On the other hand, it appears that rural birth and childhoods may moderate the effects of status threat stress and trait anxiety-depression at mPFC. Negative ruminations in trait anxiety-depression were moderated by experience of nature through mPFC function(65). It is thus conceivable that rural environments may influence stress-related mPFC development and function to being less reactive to anxiety-related traits(66, 67), as we have found, relative to urban childhoods. It remains to be fully understood if rural childhoods may contribute to more stress resilience through an inverted-U relationship that is known to scale across multiple levels of stress and physiological mechanisms(63, 68–71). It is conceivable that rural birth and childhoods bias the stress-response curve to the right of individuals with more urban childhoods, contributing to more physiological resilience, and putatively less mPFC suppression at higher levels of stress (Figure 3C, Supplementary Figure S10).

To the extent social stress-related mPFC vulnerabilities in urban vs rural childhoods affects illness-related genetic brain mechanisms, we found that these differing childhoods also influenced the effects of genome-wide genetic risk for depression. The gene-environment interaction suggest that urban childhoods potentiated not just the environmental contributions to trait anxiety-depression, but also the effects of risk-associated genetic variation for depression at the level of stress-related mPFC biology. Previous enrichment analyses of the GWAS findings to bulk tissue mRNA-seq from the Genotype-Tissue Expression (GTEx) data(72) show that the most significant enrichments were at PFC and mPFC(29). Our findings further suggest that specific aspects of urban birth and childhoods may affect genetic pathways to potentiate stress-related mPFC dysfunction. Defining these specific mechanisms should be targets of future work, and might inform potential treatment or perhaps prevention. We suggest this could involve threat-related stressors in the early environment, and their biologic interactions with genetically influenced developmental prefrontal cognitive functions.

That polygenic depression risk derived from European populations correlated with the mPFC biology of a Chinese sample supports the trans-ancestry importance of these large GWAS meta-analyses(29). We suggest at least some of these shared genetic risks are relevant at the level of biologic phenotypes influenced by critical childhood environments. We posit it is possible to leverage these diverse data to improve understanding of gene-environmental mechanisms of illness(73). Our work also extends evidence of overlapping neuropsychiatric GWAS effects across European and Chinese populations(74), including in GWAS of depression and major psychiatric disorders(75). Nevertheless, we anticipate future work on more Han Chinese-specific sets of genetic risk variation for depression, which could more powerfully define specific subsets of genetic and environmental brain mechanisms in our study population.

In conclusion, our data reflect known urban childhood advantages, such as better parental education, cognitive scores, and nutrition, but it appears there are also factors that allowed our sample of individuals with rural childhoods to cope comparably well in Beijing as adults compared with the more urban group. This appears to relate to the advantageous structural and functional effects under stress at the mPFC that we describe as potentially associated with some degree of resilience to trait anxiety-depression and polygenic risk for depression. These effects are unlikely to be confounded by genetic ancestry per se across urban-rural exposures. The more specific gene-environment mechanisms related to stress-related neuropsychiatric disorders remain to be further defined. Neural circuitry engaging the mPFC, specific urban early life exposures, and interacting neuropsychiatric risk genes would be targets for this future work.

## Acknowledgements

This work was supported by the National Natural Science Foundation of China (81361120395, D Zhang), the US National Institutes of Health (R01MH101053, Tan), the China Scholarship Council (X Zhang), and the Lieber Institute for Brain Development.

