## Supplementary materials for "Childhood urbanization affects prefrontal cortical responses to trait anxiety and interacts with polygenic risk for depression"

Supplementary Information

Supplementary Methods

### Participants

This study was approved by the Institutional Review Boards of the Peking University Institute of Mental Health and the Johns Hopkins University School of Medicine. 522 healthy subjects were initially recruited from the local community in Beijing, and written informed consent was obtained from each subject. We recruited subjects by advertising the study using social media and flyers in the community. All participants were assessed by psychiatrists using the Structured Clinical Interview for DSM-IV-TR Axis I Disorders, Research Version, Non-patient Edition (SCID-I/NP). Cognitive performance was evaluated using the MATRICS Consensus Cognitive Battery (MCCB)(1-3).

Inclusion criteria were as follows: age 18 to 45 years; right handed; Chinese of Han ancestry; no history of psychiatric or neurological diseases and substance abuse or dependence; no history of loss of consciousness duration more than 5 minutes; and no structural abnormalities on subsequent MRI, read by trained radiologists. In this study, all subjects also had to be currently living in Beijing for at least one year, and provided residence details from birth to study enrollment. They had to have at least finished the national nine-year education program, and subjects who scored 3 standard deviations below the mean in the MCCB were excluded. We excluded 2 subjects with poor quality of structural MRI images, and 490 subjects were included in the subsequent structural MRI analyses.

To determine urbanicity, subjects provided residence details from birth to present. We defined rural areas as agricultural regions with population typically <10,000; urban areas were defined as cities with populations typically more than 100,000 to well over several million. In the main text, we stratified subjects into an urban group, who had lived in cities since before they were age 12, and a rural group who were born in rural environments and only moved to cities at or after age 12. However, similar structural and functional MRI results (See Supplementary Figures S2, S3, S8) were obtained if we increased the resolution in which the differing childhood environments were quantified by stratifying subjects into 4 groups, or if we used an urbanicity score from previous studies(4, 5). In the former, the 4 groups were as follows: individuals who were born in and continue to live in cities (N=123), those who have lived in cities since before age 12 (N=126), those who lived in rural areas between birth and age 18 (N=113), and those lived in rural areas for >18 years since birth (N=128). In the case of the urbanicity score(4, 5), this was defined according to population size as follows: population < 10000 = 1, less than 1,000,000 residents = 2; cities with more than 1,000,000 residents = 3; the category scores were then multiplied by the number of years spent in the location until age 15.

### MRI Data Acquisition:

All subjects were scanned on a 3.0 T GE Discovery MR750 scanner at the Center for MRI Research, Peking University. The T1-weighted high-resolution structural image was acquired in a sagittal orientation using an axial 3D fast, spoiled gradient recalled (FSPGR) sequence with the following parameters: time repetition (TR) = 6.66 ms, time echo (TE) = 2.93 ms, field of view (FOV) = 256 × 256 mm^2^, slice thickness/gap = 1.0/0 mm, acquisition voxel size=1 × 1 × 1mm^3^, flip angle = 12°, 192 contiguous sagittal slices.

As for functional MRI, each echo-plannar image consisted of 33 (4.2 mm thick, 0 mm gap) axial slices covering the entire cerebrum and cerebellum (TR/TE = 2000/30 ms, flip angle = 90°, field of view = 22.4 cm, 64 x 64 matrix). Scanning parameters were selected to optimize the stability and quality of the BOLD signal with the exclusion of the first 4 images as dummy scans.

### Structural MRI Data Analysis

Structural MRI Data Analysis involved the following steps: (1) Transforming structural images into NIFTI format. (2) Reorienting structural images iteratively so that the millimeter coordinates of the anterior commissure (AC) matched the origin. (3) T1-weighted MR images were segmented into grey matter, white matter, cerebrospinal fluid using "New Segment" in SPM8. (4) DARTEL was used to compute transformations from individual native space to Montreal Neurological Institute (MNI) space for registration, normalization, and modulation. (5) The segmented, normalized and modulated GM images were then smoothed with an 8-mm full width at half maximum isotropic Gaussian kernel.

To study the effect of relative urban-rural childhoods on brain structure, an absolute threshold of 0.2 was used to remove voxels of low intensity from the analysis and to prevent possible edge effects(6). Voxel-based morphometry was then performed using SPM8 (<http://www.fil.ion.ucl.ac.uk/spm)>, controlling for the effects of age, second polynomial of age, gender, education years, total gray matter volume and MCCB T score(7). Significant effects were those that survived a p<0.05 whole brain family-wise error (FWE) correction. While we cannot exclude the potential effects of head motion on measures of brain structure(8), we confirmed that there were no significant group differences in the six head motion dimensions from the functional MRI acquisition performed in the same MRI session.

### Functional MRI Task and Analysis

We adapted an event-related “number working memory task” based on previous work (9, 10) (Figure 1). Subjects were trained outside the scanner for about 10 minutes. For working memory (WM), subjects encoded 2 integer numbers presented over 1s and retained in WM across a jittered interval of 3-5 seconds; in maintenance trials, subjects then responded to which of the two numbers was "larger" or "smaller" within 2s; in the manipulation trials, subjects had to perform a mental subtraction of 2 or 3 on one of the two numbers before the "larger” or “smaller” evaluation within 2s. There were 28 trials of WM manipulation and 28 trials of WM maintenance, half of which included competition and relatively more stress. Trials were embedded within equal numbers of competition or no-competition blocks, each comprising one WM maintenance and one WM manipulation trial, counterbalanced within 2 MRI runs, each about 10 minutes. Each block of competitive or non-competitive events was preceded by an initial instructional cue, whereby for competition, the participants were led to believe they were playing against a “competitor” of the same gender and of similar age, and after each WM trial, were given win or loss feedback. Here, subjects were given negative (loss) feedback approximately 70% of the time.

In the functional MRI data analysis, we excluded subjects with lower quality data according to similar criteria we have reported before (9, 10), which were accuracy rate lower than 50 percent on WM maintenance or manipulation (n = 37), those with head motion greater than 2 mm translation or 2 degrees rotation (n = 43), and those with image artifacts or did not complete the task (n = 16). Functional imaging analysis was performed using SPM12 (<http://www.fil.ion.ucl.ac.uk/spm)> and Matlab 2016b. Functional images for each subject was slice timing corrected, realigned to the first volume in the time series, and corrected for head motion. Images were then spatially normalized into standard stereotaxic space (Montreal Neurological Institute template) using a fourth degree B-spline interpolation. Spatial smoothing was applied with a Gaussian filter set at 8mm full-width at half-maximum. Each task-evoked stimulus was modeled as a separate delta function and convolved with a canonical hemodynamic response function, ratio normalized to the whole-brain global mean to control for systematic differences in global activity, and temporally filtered using a high-pass filter of 128s. Each task-evoked stimulus event was modeled for correctly performed trials. Incorrect responses and residual movement parameters were also modeled as regressors of no interest. In our study, planned contrasts of interest were brain activity at the maintenance or manipulation task phases under less stress, stress, and less stress vs stress. These contrasts were subsequently taken to a second-level analysis in which inter-subject variability was treated as a random effect.

### DNA Collection and Genotyping

Genomic DNA from the Beijing samples were extracted from peripheral blood using the QIAamp DNA Mini Kit (QIAGEN). Genotyping of samples was conducted using Illumina Human Omni ZhongHua BeadChips, designed for the Chinese population. Normalized bead intensity data obtained for each sample were loaded into Illumina BeadStudio software, which converted fluorescence intensities into SNP genotypes. Samples were excluded (N=15) according to the following quality-control criteria: (1) genotype call rate of <95%, (2) gender discordance, (3) first- or second-degree relatedness, or (4) the genetic outliers. SNPs were excluded using the following criteria: (1) minor allele frequency (MAF) <0.01, (2) genotype call rate of <95%, (3) P values for Hardy-Weinberg equilibrium < 1e-5. Principal Component Analysis (PCA) was performed to identify genetic outliers and determine whether population stratification existed between our urban and rural samples, using EIGENSTRAT (<http://genetics.med.harvard.edu/reich/Reich_Lab/Software.html)>. We compared the first 20 PCAs among the two urbanicity groups using a two-sample t-test with statistical significance set at p<0.05 corrected for the number of independent components tested.

**Supplementary Figures:**

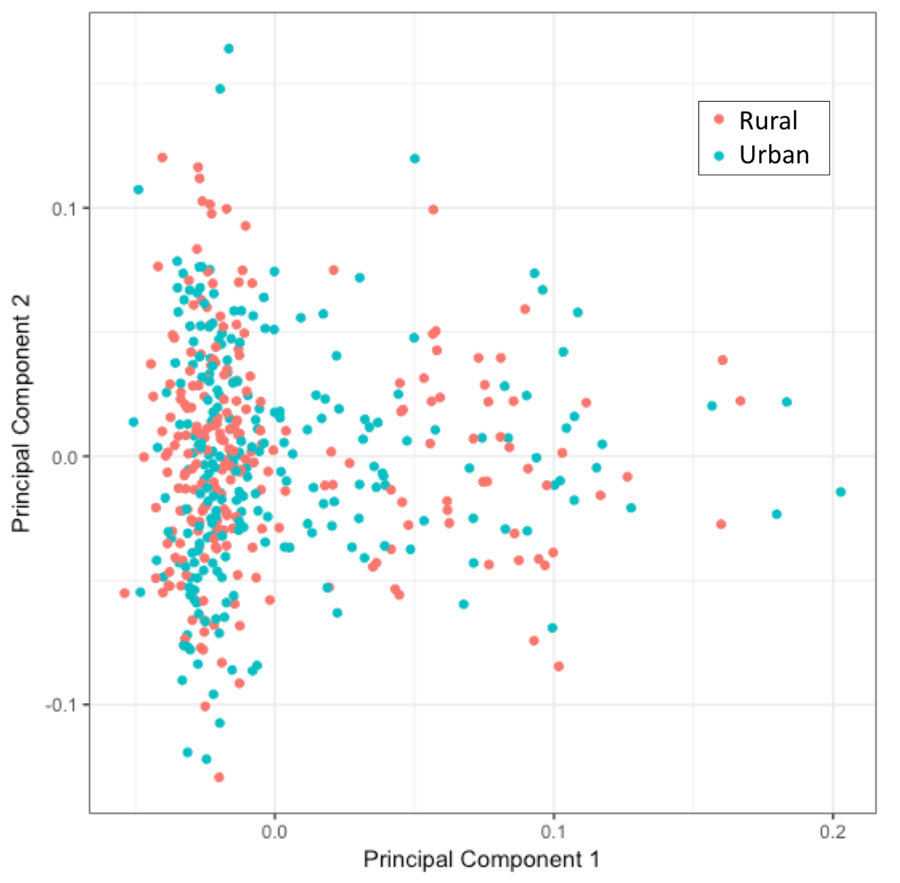

Fig. S1. *Principal components analysis on whole genome genotyping.* The plot shows overlapping first and second principal components across rural (red) and urban (blue) groups. We also compared the first 20 principal components across rural and urban groups, and there were no significant differences.

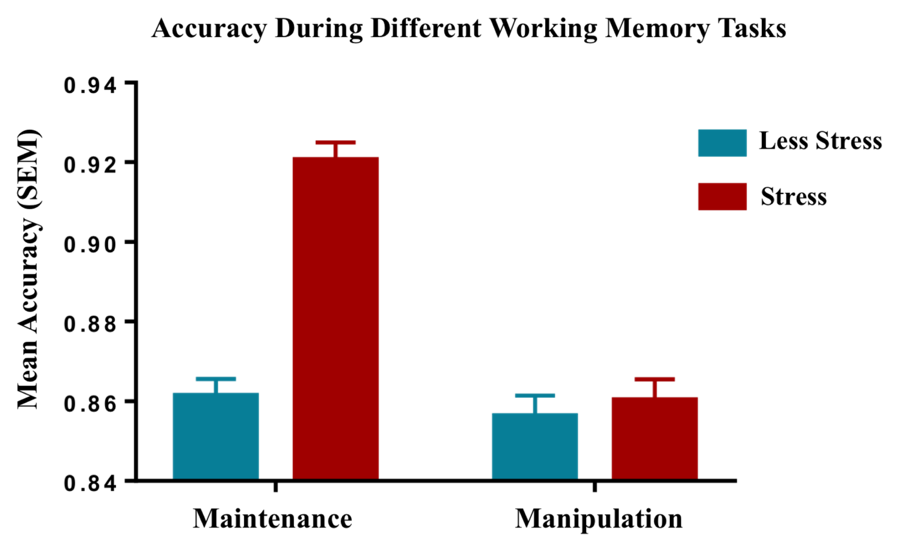

Fig. S2. *Accuracy during WM maintenance and manipulation in the whole sample (N=394).* During WM maintenance, trials with interpersonal stress were associated with relatively increased accuracy (p<0.001). This effect was not seen during WM manipulation, resulting in a significant task by stress interaction (p<0.001) consistent with a well-established bias for perseverative as opposed to flexible WM operations under stress (11, 12).

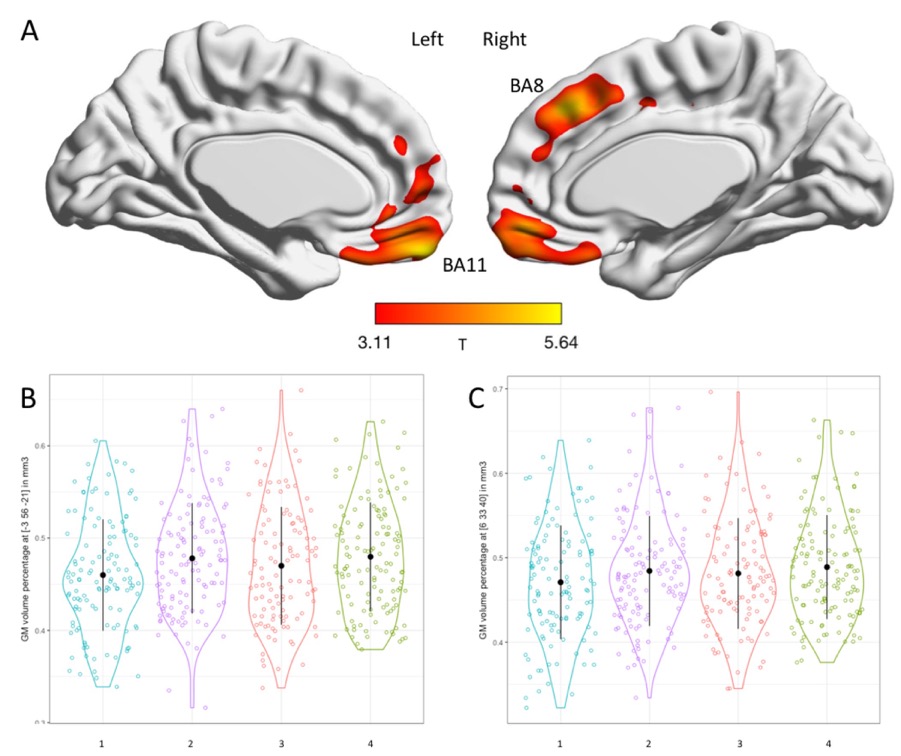

Fig. S3. *Early-life urbanicity effect on grey matter volume across four groups.* The effect of early-life urbanicity on grey matter volume was tested in a general lineal model with early-life urbanicity (Group #1: individuals who were born in and continue to live in cities; #2: individuals who have lived in cities since before age 12; #3: individuals who were born in and continue to live in rural areas until age 12-18; #4: individuals lived in rural areas for >18 years since birth). (A) T-map of the rural > urban effects controlled for age, second polynomial of age, gender, education, and total grey matter volume. (shown at p<0.001, although peaks survived p<0.05 whole-brain FWE-corrected). (B) Scatterplot of rural > urban findings at Brodmann Area 11 in medial prefrontal cortex (peak = [-3 56 -21], T = 5.64, cluster size = 556, p<0.05, whole-brain FWE-corrected). (C) Scatterplot of rural > urban findings at Brodmann Area 8 in medial prefrontal cortex (peak = [6 33 40], T = 5.48, cluster size = 198, p<0.05, whole-brain FWE-corrected).

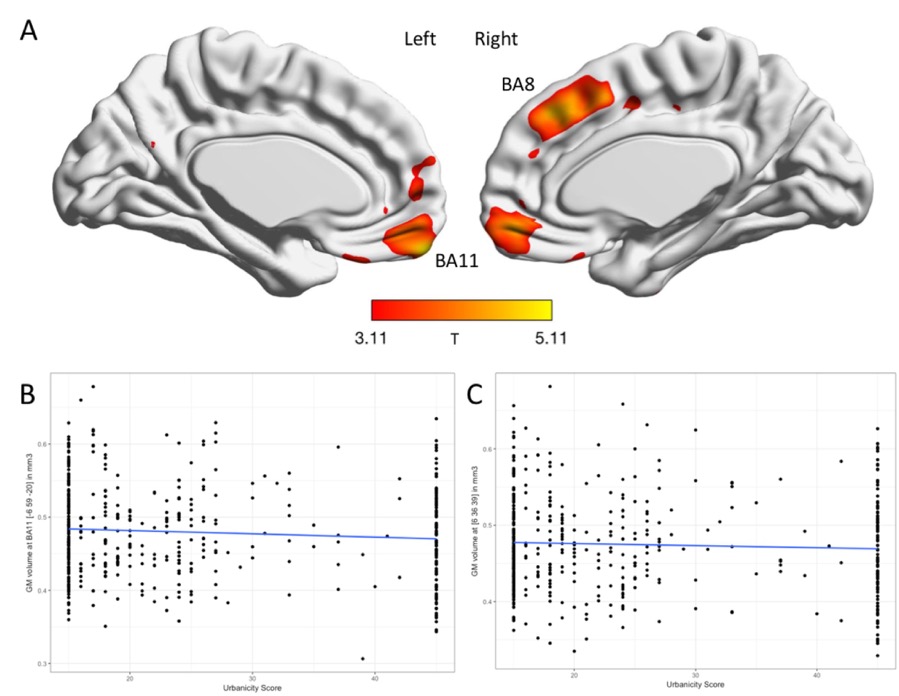

Fig. S4. *Early-life urbanicity effect on grey matter volume using the urbanicity score.* The effect of early-life urbanicity on grey matter volume was tested in a correlation analysis with early-life urbanicity score ranging from 15 to 45. (A) T-map of the rural > urban effects, controlled for age, second polynomial of age, gender, education, and total grey matter volume (shown at p<0.001 uncorrected but peaks survive p<0.05 whole-brain FWE corrected). (B) Scatterplot of rural > urban findings at Brodmann Area 11 in medial prefrontal cortex (peak = [-6 59 -20], T = 5.11, cluster size = 89, p<0.05, whole-brain FWE corrected). (C) Scatterplot of rural > urban findings at Brodmann Area 8 in medial prefrontal cortex (peak = [6 36 39], T = 4.67, cluster size = 20, p<0.05, whole-brain FWE corrected).

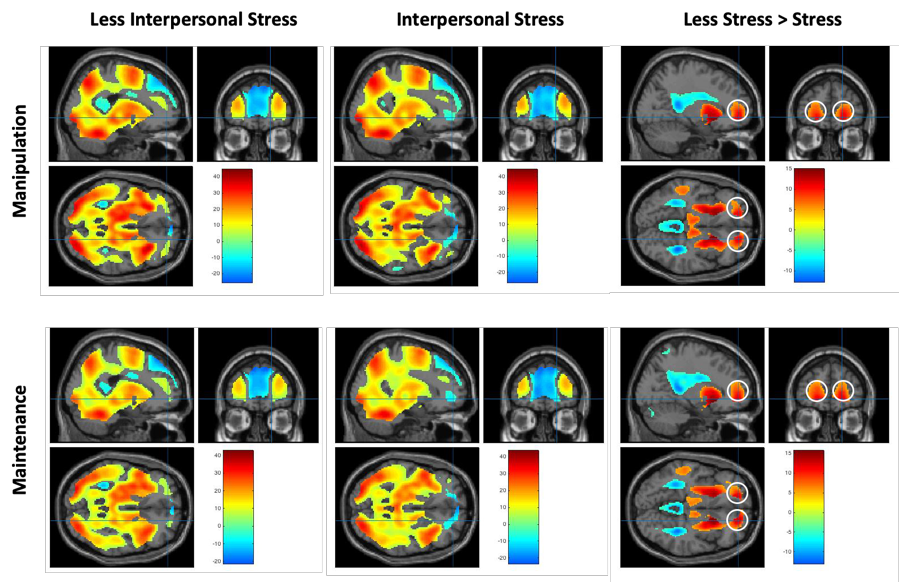

Fig. S5. *Brain activation under working memory manipulation and maintenance tasks under differing social threat stress conditions.* One-sample t-tests of working memory (WM) task across the entire sample (N=394, controlling for age; p < 0.05, whole brain FWE corrected, cluster size > 100; peaks detailed in Supplementary Tables S2-S4) under differing stress and working memory manipulation and maintenance conditions. Medial prefrontal cortex (mPFC), amongst other brain regions, was less engaged under stress. We defined 30mm diameter functional ROIs (white circles) at the mPFC sensitive to stress through the orthogonal stress vs less stress contrast during WM manipulation, or maintenance at p<0.05 whole-brain FWE-corrected. Manipulation: Right mPFC peak (ROI center), x=20, y=52, z=-6, T=11.23, 722 voxels; left peak, x=-18, y=50, z=-8, T=10.65, 542 voxels. Maintenance: Right peak, x=18, y=52, z=-6, T=12.24, 743 voxels; left peak, x=-18, y=50, z=-6, T=11.96, 725 voxels.

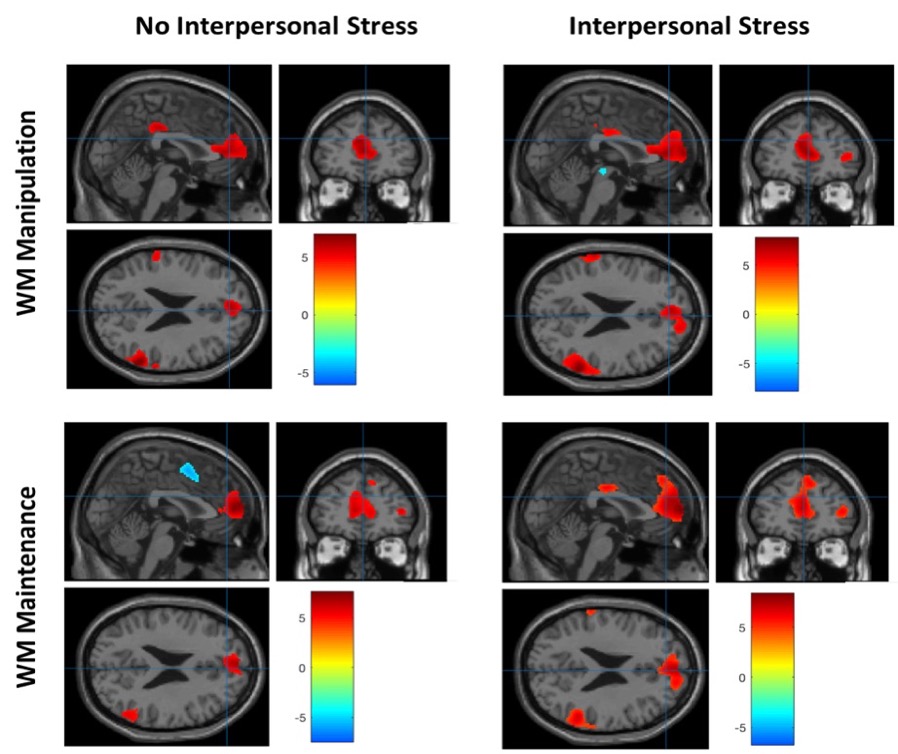

Fig. S6. *Effects of speed of processing during WM brain activation.* Correlation analysis of speed of processing (reciprocal of reaction time) and brain activation across WM manipulation and WM maintenance (N=394, controlling for age, p < 0.05 whole brain FWE corrected, cluster size > 100). Medial prefrontal cortex had enhanced engagement in relation to faster processing across WM tasks.

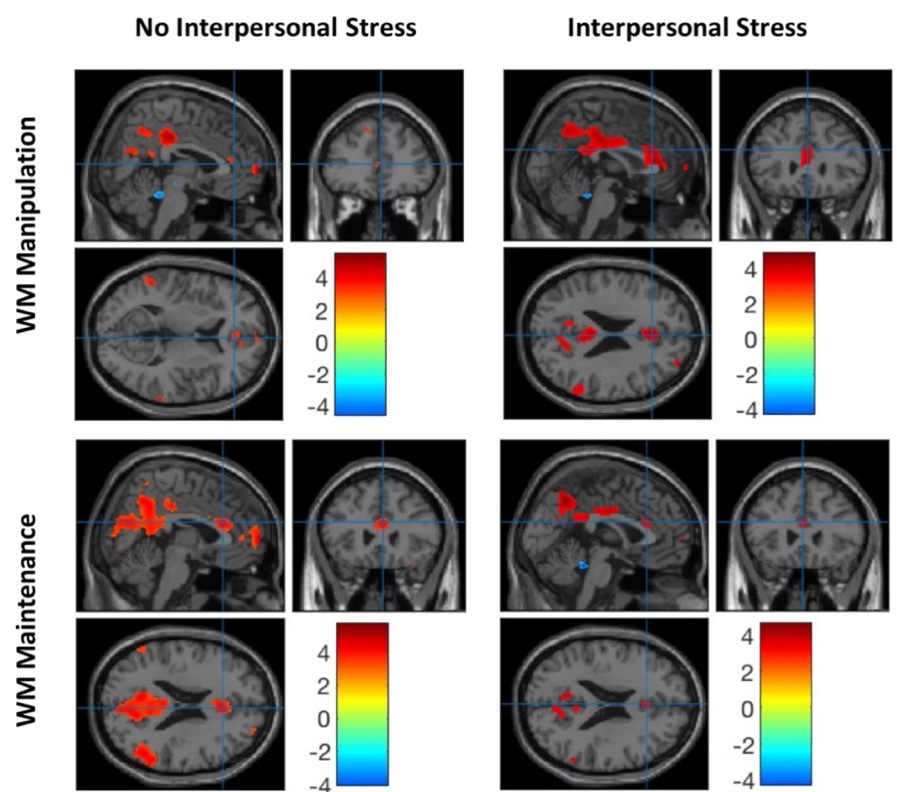

Fig. S7. *Correlation analysis of MCCB T score and brain activation.* Correlation analysis of MCCB T score and brain activation across WM manipulation and WM maintenance (N=394, controlling for age, p<0.05 FWE corrected). Medial prefrontal cortex had enhanced engagement in relation to higher overall MCCB cognitive scores across WM tasks.

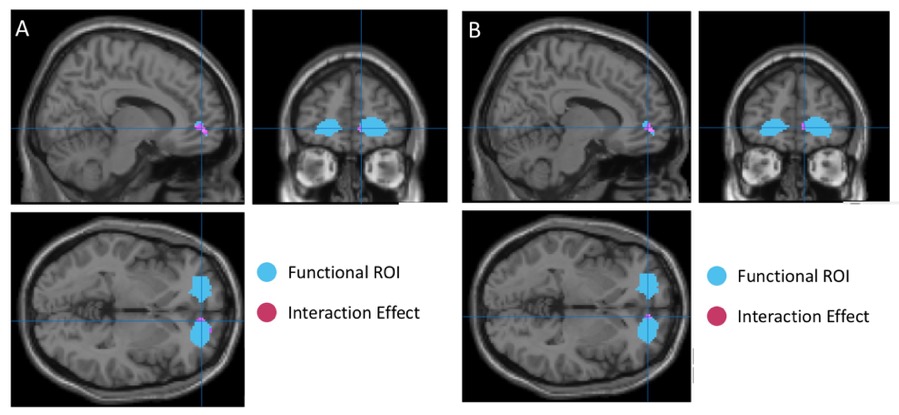

Fig. S8. *Interaction effects of trait anxiety-depression and two related definitions of urbanicity on medial prefrontal cortex engagement under interpersonal stress during working memory manipulation.* (A) Interaction effects of trait anxiety-depression and urbanicity (four categories, as in Figure S2; controlling for age and reaction time; N=394, x=8, y=54, z=-4, T=3.21; p <0.001 uncorrected). (B) Interaction effects of trait anxiety-depression and urbanicity score (Urbanicity score, as in Figure S3; controlling for age and reaction time; x=8, y=52, z=-2, T=3.19; p <0.001 uncorrected).

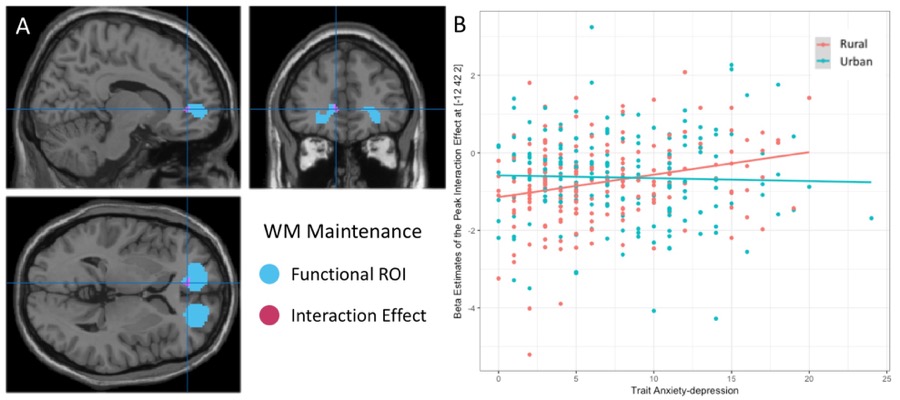

Fig. S9. *Functional effects at the mPFC ROI during working memory maintenance.* (a) Trend-level interaction effects of trait anxiety-depression and urbanicity on mPFC engagement during WM maintenance under stress (x=-12, y=42, z=2, T=2.95, controlling for age and reaction time, p<0.005 uncorrected). (b) Plot showing the interaction between rural or urban childhoods and trait anxiety-depression score.

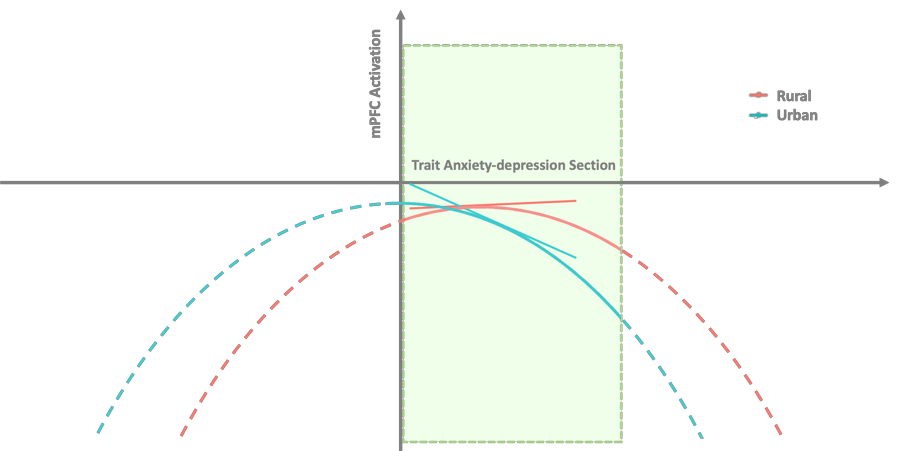

Fig. S10. *Inverted-U stress-response relationship across urbanicity.* It may be conjectured that rural birth and childhoods biased the stress-response inverted-U curve to the right of individuals with more urban childhoods, particularly during active WM manipulation under social threat stress. This may suggest a degree of physiological resilience to stress in association rural childhoods, in terms of less mPFC suppression at higher levels of stress.

Supplementary Tables:

Table S1. The effects of urbanicity, stress and task on behavioral data

| Behavior | SS | | df | MS | F | *P* |
| --- | --- | --- | --- | --- | --- | --- |
| **Accuracy** | | | | | | |
| Group (Urban/Rural) | | 0.002 | 1 | 0.002 | 0.274 | 0.600 |
| Stress (Stress/Less-Stress) | | 0.393 | 1 | 0.393 | 43.346 | < 0.001*** |
| Task (Manipulation/Maintenance) | | 0.420 | 1 | 0.420 | 46.322 | < 0.001*** |
| Group * Stress | | 0.006 | 1 | 0.006 | 0.691 | 0.406 |
| Group * Task | | 0.004 | 1 | 0.004 | 0.496 | 0.481 |
| Task*Stress | | 0.300 | 1 | 0.300 | 33.137 | < 0.001*** |
| Group * Stress * Task | | 0.001 | 1 | 0.001 | 0.148 | 0.701 |
| **Reaction Time** | | | | | | |
| Group (Urban/Rural) | 3.294 | | 1 | 3.294 | 32.050 | < 0.001*** |
| Stress (Stress/Less-Stress) | 1.262 | | 1 | 1.262 | 12.278 | < 0.001*** |
| Task (Manipulation/ Maintenance) | 61.491 | | 1 | 61.491 | 598.216 | < 0.001*** |
| Group * Stress | 0.007 | | 1 | 0.007 | 0.071 | 0.790 |
| Group * Task | 0.030 | | 1 | 0.030 | 0.290 | 0.591 |
| Stress * Task | 0.057 | | 1 | 0.057 | 0.553 | 0.457 |
| Group * Stress * Task | 0.016 | | 1 | 0.016 | 0.157 | 0.692 |

*** *P* < 0.001

Table S2. Brain activation of working memory maintenance and manipulation under social threat stress and less stress conditions (N=394, controlling for age, p< 0.05, whole-brain FWE corrected, cluster size > 100).

| Peak Region | Cluster | x | y | z | T score |
| --- | --- | --- | --- | --- | --- |
| **Manipulation: Stress** | | | | | |
| L Middle Frontal Gyrus | 112370 | -32 | 50 | 16 | 18.74 |
| R Middle Frontal Gyrus |  | 44 | 36 | 24 | 25.30 |
| R Cingulate Gyrus |  | 6 | 16 | 44 | 41.27 |
| L Middle Frontal Gyrus |  | -44 | 20 | 28 | 29.35 |
| R Middle Frontal Gyrus |  | 54 | 12 | 34 | 30.27 |
| R Insula |  | 32 | 24 | 0 | 34.69 |
| L Insula |  | -34 | 18 | 2 | 33.14 |
| L Insula |  | -40 | -2 | 6 | 37.20 |
| L Thalamus |  | -18 | -10 | -2 | 30.00 |
| R Thalamus |  | 14 | -12 | 10 | 27.41 |
| L Inferior Parietal Lobule |  | -44 | -32 | 44 | 44.23 |
| R Inferior Parietal Lobule |  | 46 | -36 | 42 | 33.00 |
| L Hippocampus |  | -20 | -30 | -6 | 27.95 |
| R Hippocampus |  | 24 | -26 | -10 | 24.37 |
| R Culmen |  | 32 | -54 | -32 | 42.33 |
| R Middle Occipital Gyrus |  | 30 | -68 | 32 | 33.09 |
| L Precuneus |  | -24 | -66 | 40 | 42.01 |
| L Middle Occipital Gyrus |  | -28 | -88 | -2 | 37.57 |
| R Occipital Lobe |  | 28 | -86 | -6 | 37.60 |
| **Manipulation: Less Stress** | | | | | |
| L Inferior Parietal Lobule | 108004 | -46 | -34 | 44 | 44.11 |
| R Cingulate Gyrus |  | 6 | 16 | 42 | 43.83 |
| L Middle Frontal Gyrus |  | -30 | 48 | 14 | 21.06 |
| R Middle Frontal Gyrus |  | 32 | 52 | 16 | 19.33 |
| R Middle Frontal Gyrus |  | 44 | 34 | 26 | 26.48 |
| L Middle Frontal Gyrus |  | -44 | 20 | 26 | 31.29 |
| R Inferior Frontal Gyrus |  | 52 | 12 | 34 | 33.11 |
| L Inferior Frontal Gyrus |  | -52 | 10 | 26 | 36.59 |
| R Insula |  | 32 | 24 | -2 | 38.33 |
| L Insula |  | -30 | 20 | 2 | 37.32 |
| L Medial Frontal Gyrus |  | -2 | 6 | 50 | 42.42 |
| L Middle Frontal Gyrus |  | -30 | -4 | 56 | 41.90 |
| R Middle Frontal Gyrus |  | 28 | 0 | 58 | 31.43 |
| L Lentiform Nucleus |  | -20 | -6 | 0 | 32.68 |
| R Lentiform Nucleus |  | 16 | -4 | -2 | 28.63 |
| L Insula |  | -40 | -4 | 8 | 36.05 |
| R Insula |  | 42 | 0 | 6 | 27.88 |
| L Thalamus |  | -12 | -20 | 4 | 37.64 |
| R Thalamus |  | 12 | -14 | 6 | 29.80 |
| R Superior Parietal Lobule |  | 26 | -68 | 46 | 35.92 |
| L Precuneus |  | -26 | -64 | 42 | 43.71 |
| L Middle Occipital Gyrus |  | -30 | -88 | -2 | 35.98 |
| R Inferior Occipital Gyrus |  | 28 | -86 | -8 | 35.68 |
| **Maintenance: Stress** | | | | | |
| L Middle Frontal Gyrus | 115766 | -40 | 34 | 24 | 22.09 |
| R Middle Frontal Gyrus |  | 40 | 40 | 28 | 22.62 |
| R Cingulate Gyrus |  | 6 | 16 | 40 | 38.60 |
| R Insula |  | 34 | 20 | 2 | 29.90 |
| L Insula |  | -32 | 18 | 4 | 29.60 |
| L Insula |  | -40 | -2 | 6 | 40.85 |
| R Insula |  | 46 | 4 | 2 | 30.92 |
| L Thalamus |  | -12 | -20 | 6 | 27.49 |
| R Thalamus |  | 6 | -20 | 8 | 25.67 |
| L Inferior Parietal Lobule |  | -44 | -32 | 44 | 43.31 |
| R Postcentral Gyrus |  | 48 | -30 | 40 | 26.36 |
| L Precuneus |  | -18 | -70 | 48 | 34.21 |
| R Inferior Parietal Lobule |  | 32 | -58 | 48 | 31.64 |
| L Middle Occipital Gyrus |  | -26 | -88 | -2 | 32.78 |
| R Middle Occipital Gyrus |  | 28 | -86 | -4 | 31.55 |
| L Hippocampus |  | -18 | -30 | -8 | 23.30 |
| **Maintenance: Less Stress** | | | | | |
| L Middle Frontal Gyrus | 110896 | -32 | 46 | 14 | 18.25 |
| R Middle Frontal Gyrus |  | 30 | 46 | 22 | 19.61 |
| L Inferior Frontal Gyrus |  | -56 | 8 | 26 | 34.03 |
| R Inferior Frontal Gyrus |  | 54 | 10 | 36 | 30.39 |
| Medial Frontal Gyrus |  | 0 | 4 | 50 | 36.18 |
| L Insula |  | -30 | 18 | 2 | 31.40 |
| R Insula |  | 32 | 20 | 2 | 29.56 |
| L Insula |  | -40 | -2 | 6 | 37.76 |
| R Insula |  | 40 | 0 | 6 | 29.47 |
| L Thalamus |  | -12 | -20 | 4 | 30.62 |
| R Thalamus |  | 14 | -14 | 4 | 22.84 |
| L Postcentral Gyrus |  | -38 | -28 | 52 | 37.50 |
| R Postcentral Gyrus |  | 50 | -24 | 38 | 25.86 |
| L Precuneus |  | -26 | -64 | 42 | 36.75 |
| R Inferior Parietal Lobule |  | 32 | -58 | 48 | 30.93 |
| L Middle Occipital Gyrus |  | -28 | -88 | -2 | 32.43 |
| R Occipital Lobe |  | 28 | -86 | -6 | 29.41 |

Table S3. Activation contrasts across working memory maintenance and manipulation under stress and less stressed conditions (N=394, controlling for age, p< 0.05, whole-brain FWE-corrected, cluster size > 100).

| Peak Region | Cluster | x | y | z | T score |
| --- | --- | --- | --- | --- | --- |
| **Stress: Manipulation > Maintenance** | | | | | |
| R Insula | 65620 | 32 | 26 | 0 | 19.54 |
| L Insula |  | -30 | 26 | 0 | 19.61 |
| L Middle Frontal Gyrus |  | -30 | -2 | 56 | 18.26 |
| L Medial Frontal Gyrus |  | -2 | 12 | 52 | 23.56 |
| R Inferior Frontal Gyrus |  | 46 | 8 | 26 | 19.56 |
| R Middle Frontal Gyrus |  | 28 | -2 | 50 | 19.20 |
| L Thalamus |  | -12 | -14 | 4 | 14.5 |
| R Thalamus |  | 12 | -16 | 8 | 11.07 |
| L Hippocampus |  | -22 | -32 | -6 | 13.87 |
| R Hippocampus |  | 24 | -28 | -8 | 12.13 |
| L Inferior Parietal Lobule |  | -46 | -40 | 42 | 19.51 |
| R Inferior Parietal Lobule |  | 46 | -38 | 42 | 19.65 |
| R Superior Parietal Lobule |  | 26 | -66 | 44 | 28.04 |
| L Precuneus |  | -24 | -68 | 42 | 26.03 |
| L Middle Occipital Gyrus |  | -30 | -82 | 10 | 26.75 |
| R Middle Occipital Gyrus |  | 34 | -82 | 2 | 28.22 |
| **Less Stress: Manipulation > Maintenance** | | | | | |
| L Insula | 64008 | -30 | 26 | 0 | 20.29 |
| R Insula |  | 32 | 26 | -2 | 19.93 |
| L Inferior Frontal Gyrus |  | -44 | 6 | 28 | 26.71 |
| R Inferior Frontal Gyrus |  | 48 | 10 | 26 | 21.57 |
| L Medial Frontal Gyrus |  | -4 | 12 | 50 | 25.33 |
| L Thalamus |  | -12 | -14 | 0 | 16.65 |
| R Thalamus |  | 12 | -18 | 8 | 12.47 |
| R Middle Frontal Gyrus |  | 28 | 0 | 54 | 20.15 |
| L Middle Frontal Gyrus |  | -28 | -2 | 52 | 19.75 |
| R Inferior Parietal Lobule |  | 48 | -38 | 46 | 22.61 |
| L Inferior Parietal Lobule |  | -44 | -42 | 42 | 20.72 |
| L Precuneus |  | -24 | -68 | 38 | 27.52 |
| R Precuneus |  | 28 | -68 | 42 | 29.38 |
| L Middle Occipital Gyrus |  | -30 | -84 | 6 | 24.61 |
| R Middle Occipital Gyrus |  | 36 | -82 | 2 | 28.54 |
| **Stress:** **Maintenance > Manipulation** | | | | | |
| L Cingulate Gyrus | 4666 | -4 | -42 | 34 | 15.60 |
| R Cingulate Gyrus |  | 10 | -48 | 30 | 15.53 |
| L Precuneus |  | -10 | -64 | 22 | 10.88 |
| L Angular Gyrus | 1666 | -46 | -74 | 34 | 15.48 |
| L Superior Temporal Gyrus |  | -56 | -64 | 28 | 11.92 |
| L Inferior Parietal Lobule |  | -66 | -36 | 28 | 9.54 |
| R Angular Gyrus | 5762 | 52 | -68 | 32 | 15.48 |
| R Middle Temporal Gyrus |  | 62 | -40 | -2 | 13.75 |
| R Superior Temporal Gyrus |  | 60 | -58 | 16 | 13.52 |
| R Inferior Parietal Lobule |  | 62 | -26 | 26 | 12.14 |
| R Middle Temporal Gyrus |  | 52 | -8 | -14 | 10.17 |
| R Superior Frontal Gyrus | 2168 | 14 | 50 | 36 | 8.85 |
| R Middle Frontal Gyrus |  | 24 | 30 | 46 | 8.04 |
| L Medial Frontal Gyrus |  | -6 | 62 | 4 | 6.86 |
| L Middle Temporal Gyrus | 896 | -58 | -14 | -14 | 8.25 |
| L Insula |  | -42 | -12 | -6 | 7.90 |
| L Middle Temporal Gyrus |  | -54 | -2 | -18 | 7.03 |
| L Middle Frontal Gyrus | 273 | -24 | 26 | 42 | 6.89 |
| L Superior Frontal Gyrus |  | -16 | 46 | 34 | 5.19 |
| **Less Stress: Maintenance > Manipulation** | | | | | |
| R Cingulate Gyrus | 4728 | 8 | -50 | 28 | 16.84 |
| L Cingulate Gyrus |  | -4 | -44 | 32 | 15.52 |
| L Precuneus |  | -10 | -64 | 20 | 12.66 |
| L Angular Gyrus | 1793 | -46 | -74 | 34 | 15.12 |
| L Superior Temporal Gyrus |  | -58 | -62 | 22 | 9.56 |
| L Supramarginal Gyrus |  | -60 | -52 | 30 | 8.76 |
| R Superior Temporal Gyrus | 6236 | 62 | -54 | 10 | 14.53 |
| R Postcentral Gyrus |  | 62 | -26 | 22 | 13.77 |
| R Middle Temporal Gyrus |  | 60 | -40 | -2 | 13.21 |
| R Superior Frontal Gyrus | 3546 | 4 | 60 | -6 | 10.86 |
| R Superior Frontal Gyrus |  | 14 | 50 | 38 | 9.60 |
| R Medial Frontal Gyrus |  | 8 | 56 | 14 | 8.56 |
| L Insula | 1121 | -42 | -12 | -8 | 8.18 |
| L Insula |  | -42 | -18 | -2 | 7.80 |
| L Middle Temporal Gyrus |  | -62 | -14 | -14 | 7.55 |
| R Inferior Frontal Gyrus | 106 | 52 | 32 | 0 | 7.83 |

Table S4. Brain activation contrasts between stress and less stress conditions during working memory maintenance or manipulation (N=394, controlling for age, p< 0.05, whole-brain FWE-corrected, cluster size > 100).

| Peak Region | Cluster | x | y | z | T score |
| --- | --- | --- | --- | --- | --- |
| **Manipulation: Less Stress > Stress** | | | | | |
| R Putamen | 3627 | 18 | 14 | -8 | 14.82 |
| R Medial Frontal Lobe |  | 20 | 52 | -6 | 11.23 |
| R Amygdala |  | 28 | -8 | -12 | 11.19 |
| L Extra-Nuclear | 3657 | -22 | 16 | -10 | 14.67 |
| L Putamen |  | -22 | 4 | -6 | 13.05 |
| L Caudate |  | -12 | 14 | 4 | 11.04 |
| L Inferior Frontal Lobe | 2531 | -40 | 8 | 28 | 9.04 |
| L Inferior Frontal Gyrus |  | -44 | 2 | 36 | 8.11 |
| R Middle Frontal Gyrus | 1667 | 46 | 12 | 30 | 8.85 |
| R Middle Frontal Gyrus |  | 54 | 26 | 32 | 7.57 |
| R Middle Frontal Gyrus |  | 44 | 30 | 22 | 6.45 |
| R Inferior Parietal Lobule | 1056 | 38 | -62 | 46 | 8.79 |
| R Midbrain | 933 | 10 | -18 | -6 | 8.60 |
| L Midbrain |  | -8 | -18 | -8 | 8.10 |
| L Thalamus |  | -16 | -22 | 0 | 6.84 |
| L Middle Temporal Gyrus | 285 | -51 | -32 | -6 | 7.02 |
| L Precuneus | 538 | -30 | -64 | 38 | 6.89 |
| **Maintenance: Less Stress > Stress** | | | | | |
| R Putamen | 4075 | 18 | 14 | -8 | 15.52 |
| R Medial Frontal Lobe |  | 18 | 52 | -6 | 12.44 |
| R Superior Temporal Gyrus |  | 42 | 18 | -36 | 8.52 |
| L Putamen | 4117 | -20 | 14 | -8 | 13.96 |
| L Frontal Lobe |  | -18 | 50 | -6 | 11.96 |
| L Anterior Cingulate |  | -10 | 40 | 2 | 9.28 |
| R Precentral Gyrus | 840 | 58 | -6 | 46 | 6.97 |
| R Precentral Gyrus |  | 54 | -4 | 28 | 6.27 |
| R Precentral Gyrus |  | 52 | -12 | 54 | 5.66 |
| L Brain Stem | 314 | -10 | -12 | -10 | 6.87 |
| L Middle Temporal Gyrus | 416 | -58 | -30 | -4 | 6.8 |
| L Middle Temporal Gyrus |  | -52 | -40 | 0 | 6.7 |
| L Middle Temporal Gyrus |  | -56 | -16 | -8 | 4.93 |
| L Middle Frontal Gyrus | 1397 | -42 | 0 | 44 | 6.72 |
| L Middle Frontal Gyrus |  | -42 | 10 | 30 | 6.14 |
| **Manipulation: Stress > Less Stress** | | | | | |
| R Fusiform | 12046 | 34 | -42 | -4 | 12.54 |
| L Temporal Lobe |  | -34 | -46 | -2 | 10.35 |
| L Lateral Ventricle |  | -18 | -44 | 14 | 9.70 |
| R Lateral Ventricle |  | 20 | -42 | 14 | 11.34 |
| R Extra-Nuclear |  | 20 | -2 | 26 | 8.17 |
| L Extra-Nuclear |  | -18 | -2 | 26 | 9.99 |
| Inter-Hemispheric |  | 0 | -26 | 6 | 10.09 |
| L Lateral Ventricle |  | -28 | -52 | 10 | 11.21 |
| R Lateral Ventricle |  | 18 | -40 | 12 | 11.47 |
| L Precuneus | 898 | -10 | -60 | 66 | 8.99 |
| R Precuneus |  | 6 | -60 | 66 | 7.45 |
| **Maintenance: Stress > Less Stress** | | | | | |
| L Extra-Nuclear | 12370 | -20 | -4 | 26 | 10.16 |
| R Extra-Nuclear |  | 20 | -6 | 26 | 9.28 |
| L Extra-Nuclear |  | -22 | -16 | 26 | 10.97 |
| R Extra-Nuclear |  | 26 | -44 | 18 | 11.19 |
| L Lateral Ventricle |  | -22 | -36 | 12 | 10.20 |
| L Para-Hippocampus |  | -34 | -44 | -4 | 11.12 |
| R Lateral Ventricle |  | 34 | -42 | -2 | 12.30 |
| Inter-Hemispheric |  | 0 | -24 | 8 | 12.80 |
| R Lateral Ventricle |  | 20 | -40 | 10 | 11.62 |
| R Postcentral Gyrus | 957 | 8 | -58 | 68 | 8.13 |
| R Precuneus |  | 8 | -74 | 54 | 7.76 |
| R Precuneus |  | 6 | -68 | 60 | 7.47 |
| L Inferior Parietal Lobule | 105 | -66 | -30 | 30 | 5.96 |
